# Dynamic interplay between a TonB dependent heme transporter and a TonB protein in a membrane environment

**DOI:** 10.1101/2021.04.21.440789

**Authors:** Kamolrat Somboon, Oliver Melling, Maylis Lejeune, Glaucia M.S. Pinheiro, Annick Paquelin, Benjamin Bardiaux, Michael Nilges, Philippe Delepelaire, Syma Khalid, Nadia Izadi-Pruneyre

**Affiliations:** School of Chemistry, University of Southampton, Southampton S017 1BJ, United Kingdom; Institut Pasteur, Université Paris Cité, CNRS UMR3528, Structural Bioinformatics Unit, F-75015 Paris, France; Institut de Biologie Physico-Chimique, UMR 7099, CNRS Université de Paris, 13 rue Pierre et Marie Curie, Paris, France; Department of Biochemistry, University of Oxford, Oxford, OX1 3QU, United Kingdom

**Author notes:** Corresponding authors: Syma Khalid & Nadia Izadi-Pruneyre, **Email:**.

## Abstract

The envelope of Gram-negative bacteria is composed of a double membrane separated by the periplasmic space. This organization imposes geometrical and distance constraints that are key for the mechanism of action of multicomponent systems spanning the envelope. However, consideration of all three compartments by experimental approaches is still elusive. Here we used the state-of-the-art molecular dynamics simulation in an *Escherichia coli* envelope model to obtain a dynamic view of molecular interactions between the outer membrane heme transporter HasR and the inner membrane TonB-like protein HasB. Their interaction allows the transfer of the inner membrane proton motive force derived energy to the transporter for heme internalization. The simulations which incorporate both membranes show the key role of periplasmic domains of both proteins, and their dynamics in the complex formation and stability. They revealed a previously unidentified atomic network of interactions, as well as the sequences of the interactions and their variations with the presence of external substrates. Experimental validation (mutations, phenotypic and *in vitro* assays) confirms the robustness of our approach and provides verification of the simulation-predicted interactions. Based on structural and sequence conservation, the network of interaction revealed in this study is expected to occur in other nutrient import systems. The integrative approach presented here is highly useful to study the dynamic interplay between components of any other bacterial transmembrane systems.

## Introduction

Gram-negative bacteria are delimited by an envelope composed of two membranes (outer membrane, OM and inner membrane, IM) separated by the periplasmic space. The IM TonB complex and the specific OM TonB-dependent transporters (TBDTs) are used to import scarce and essential nutrients, such as metal sources, vitamin B_12_, some specific carbohydrates, etc (1, 2). As both the OM and the periplasm lack an energy source, the TonB complex is used to couple substrate entry to the proton-motive force provided by the proton gradient across the IM.

The TonB complex comprises three IM proteins, TonB, ExbB and ExbD. TonB is the only protein of the complex interacting directly with the TBDT and thus transferring the energy, whereas ExbB and ExbD, forming a proton-channel, are considered as the engine of the system (3). The structure of the complete TonB complex and of the full-length TonB protein are still unknown. TonB is composed of a N-terminus transmembrane helix inserted into the IM, an unstructured proline-rich domain in the periplasm followed by a C-terminal and globular domain that is the only identified domain of interaction with TBDT. Only the structure of this globular periplasmic domain of TonB, either alone or in complex with TBDT, have been reported (4–8). TonB interacts with the TonB-box, a stretch of 7-10 conserved residues, exposed in the periplasm and located at the N-terminus of TBDT. It is assumed that upon this interaction and in the presence of energy, by either a rotation (comparable to the Mot flagellar system)(9) or a linear movement of TonB, some conformational changes of the TBDT occur, eventually opening a channel for the entry of the substrate. This energising mechanism is not well understood yet due to the lack of structural and interaction data on the whole system.

Here we use the heme acquisition system (Has) to get insight into the dynamic interplay between a TBDT and a TonB protein. The Has is developed by commensal and pathogenic bacteria to acquire heme that is their major iron source in mammals. The system studied here is from *Serratia marcescens* and reconstituted in *Escherichia coli*. It is composed of the TBDT HasR functioning in synergy with a hemophore (HasA) that harvests extracellular heme, either free or bound to host hemoproteins (like hemoglobin). Heme is then transferred from HasA to HasR to be internalized, the complex formed by HasA-HasR is dissociated, and HasA is recycled into the external medium. The two steps of heme internalization and recycling of HasA need energy that is provided by HasB, a TonB paralog dedicated to the Has system(10–12).

HasR has the same structural organization as other TBDTs, with a 22-stranded β-barrel forming a trans-membrane domain anchored in the OM, with 11 extracellular loops and 11 periplasmic loops. The region upstream this domain folds inside the barrel, forming the so-called “plug”(1). HasR belongs to a class of TBDTs coupled to a signalling activity. In these TBDTs, the TonB-box is not at the extreme N-terminus, but is connected to a periplasmic globular domain called the signalling domain, or HasR_SD_ in HasR(13). HasR_SD_ sends the signal of the presence of extracellular sources of heme (heme and HasA) to an anti-sigma/ECFσ factor pair which activates the expression of the *has* operon (10).

Previously we combined the X-ray structure of HasR (14), the NMR structure of HasR_SD_ with electron microscopy (EM) density maps and SAXS data, to obtain a structural model of the full-length HasR in its free form and in complex with HasA and heme (holo-HasR). This model showed for the first time the structure of a full-length TBDT, with the signalling domain projected far from the β-barrel (at 8 and 7 nm in the apo- and holo-form, respectively)(13) *via* a flexible linker. A similar location of the signalling domain with respect to the β-barrel was recently shown in the X-ray structure of FoxA from *Pseudomonas aeruginosa*(8).

HasR and HasB interact through their periplasmic exposed regions. In previous work, we solved the structure of the C-terminal and periplasmic domains of HasB (HasB_CTD_)(15) and studied its interaction with a peptide containing the TonB-box of HasR (G_95_ALALDSL_102_). We showed that an ionic interaction between HasR (D_100_) and HasB (R_169_) is crucial for the stability of this interaction and for HasR function. Furthermore, by using isothermal titration microcalorimetry (ITC), we showed that heme and HasA modulate the thermodynamic parameters and the network of interactions(16). However, the precise nature of the interactions, their variations and the interacting residues remained unknown.

HasR and HasB are inserted into two different membranes that are separated by the periplasmic space. To characterise the HasR-HasB interaction, it is therefore critical to consider the spatial constraints imposed by the membranes, in particular the distance between them together with the protein dynamics. In this work, to develop a more thorough view of the HasR-HasB interaction, we have combined our structural and interaction data to model the protein complex in the context of *in silico* models of the IM and OM. We performed molecular dynamics (MD) simulations of the HasR+HasB system in apo- and holo-forms, that is to say in the presence of heme and HasA (holoHasR+HasB). The simulations revealed a previously unidentified network of interactions and the key role of particular periplasmic loops in the complex formation and stability. We experimentally validated the network of interactions by mutations, and phenotypic and *in vitro* assays. Interestingly, the interactions are modulated by the external signal consisting in the presence of heme and HasA. This work represents the first example showing the molecular details of the interaction of a TBDT with a TonB like protein with inclusion of both membranes. The sequence of events and the movements observed during the simulations provide a more detailed understanding of the mechanism of energy and signal transfer through the bacterial cell envelope.

## Results

### Molecular Dynamics of the HasR-HasB complex

We started by the simplest system (HasR-HasB) consisting of HasR in apo-form (without its extracellular substrates) and HasB (Figure 1). We constructed an initial model of these two proteins each in a membrane model based on their structural organization, and guided by our previous experimental findings, as described in the supplementary Information. In the HasR model, the linker was almost fully extended at a distance of 11.5 nm from the barrel. The HasB_CTD_ is located at ∼ 6.8 nm from the HasR barrel (centre of mass measurements). The HasB transmembrane domain is a single helix located in the model IM. The two membranes are separated by ∼ 12 nm, this is the shortest distance (the distance from the centre of the OM to the centre of the IM is ∼ 16 nm). This was deemed to be a good compromise between the average of the periplasmic thicknesses reported in the literature and keeping the simulation system size practical. (Figure 1)

**Figure 1.**
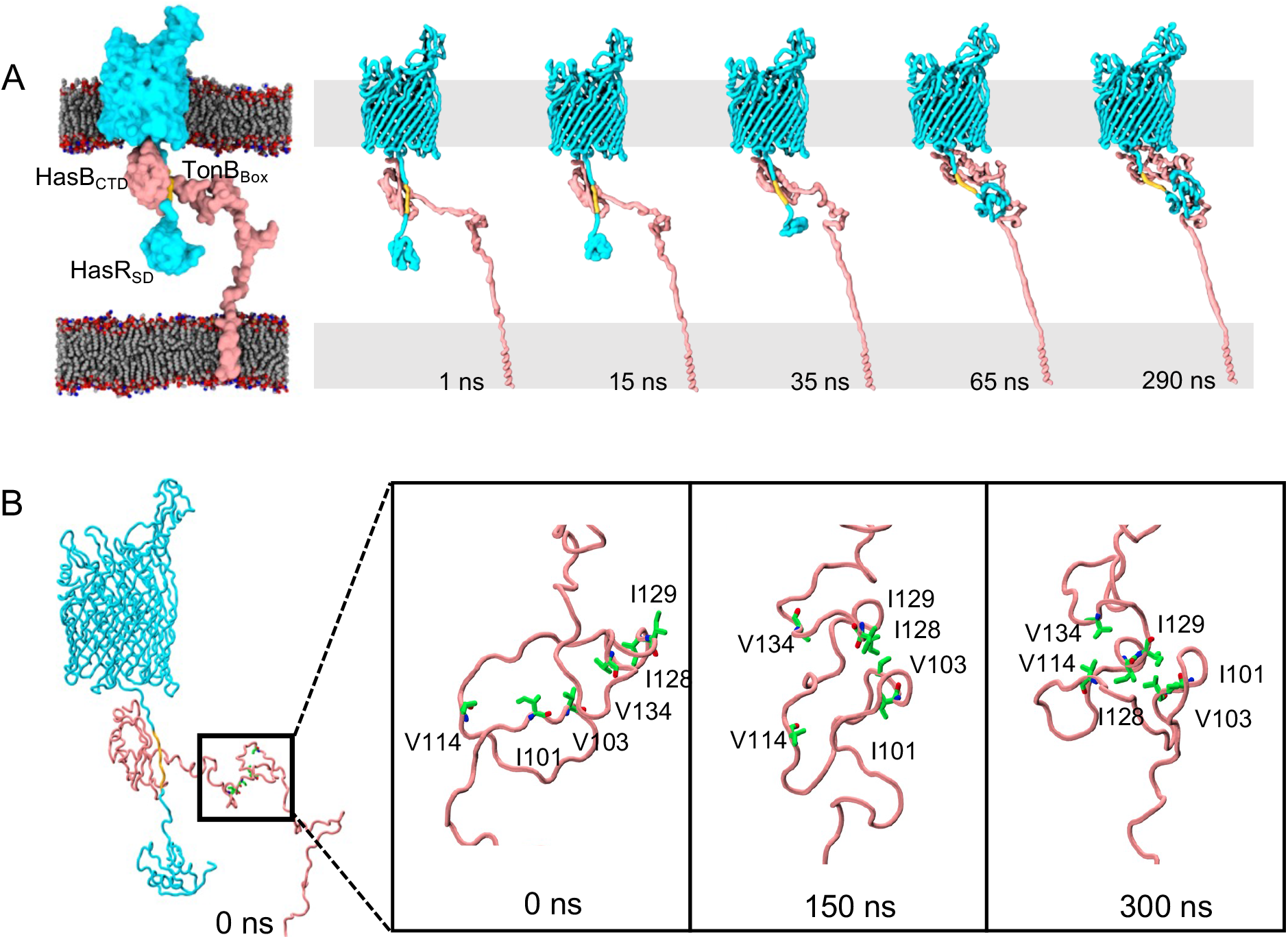
The HasR-HasB complex. **A. PCA analysis**. HasR is cyan and HasB is pink. Left: Initial model of the HasR-HasB complex at time = 0 ns. Each protein is embedded in a model membrane. The membrane lipids are grey, red and blue, the TonB box is in yellow, water and ions are omitted for clarity. Proteins are shown in surface representation, while membrane is shown as spheres. Right: Spontaneous interaction between HasB and HasR periplasmic loops. Here only the protein backbones are included in the surface representation. Projection along the first principle component from the HasR-HasB simulation at 1, 15, 35, 65 and 290 ns. **B. Hydrophobic contacts along the HasB proline-rich domain**. The panel on the left shows the location of the hydrophobic residues at 0 ns. The close-up version on the right shows this domain becoming less extended during the simulation. Key hydrophobic residues in HasB are shown as sticks and labelled.

### Visual observations

On performing two initial simulations of the HasR-HasB model as described above, in one of the simulations we noted interaction of two periplasmic loops of HasR with HasB_CTD_ (discussed in detail later), so we initiated two additional simulations of the HasR-HasB complex from a point at which the contact between the loops of HasR and HasB has already been established (Figure 1). Thus, overall we simulated the HasR-HasB complex as (1) HasR-HasB in which wild-types of both proteins are considered in the original model (2 × 300 ns), (2) int-HasR-HasB in which the simulations are initiated after the contact with the loops has been made (4 × 300 ns), and (3) HasR-HasB with a periplasmic space at 23 nm called the wide periplasm hereafter (4 × 50 ns) (4) HasR_mutants_-HasB in which different periplasmic loops of HasR are mutated as described below (2 × 300 ns per mutant).

### Secondary structure and structural integrity

Given that the starting conformations of the complexes we are simulating are models, albeit based on structural data and guided by experimental data, we first evaluated the structural integrity of the folded domains. The root mean square deviation (RMSD) from the initial model and the secondary structures were evaluated. In all cases, we observed that the protein structural deviation is within acceptable ranges based on simulations of similar proteins and there is no significant unfolding of the structured domains. The RMSD, root mean square fluctuation (RMSF) and secondary structure analysis of one simulation of each system are provided in the supplementary information (Figure S1).

### Inter-protein motion

The long disordered periplasmic regions of HasR and HasB impart considerable flexibility to the complex in the periplasm. Therefore, we next explored the inter-domain and inter-protein movements. A useful way of filtering the large-scale motions is principal components analysis (PCA). We employed this method to characterise the motion along the first eigenvector for the HasR-HasB complexes in the HasR-HasB and int-HasR-HasB systems (Figure 1). Interpolation between the extreme conformations sampled in each case shows a distinct pattern in the inter-protein motions. The principle motion in the original HasR-HasB simulations is a reduction in the distance between the HasR_SD_ and the barrel, and a concomitant reduction in the distance between HasB_CTD_ and HasR_SD_ as shown in Figure 1 and 2.

**Figure 2.**
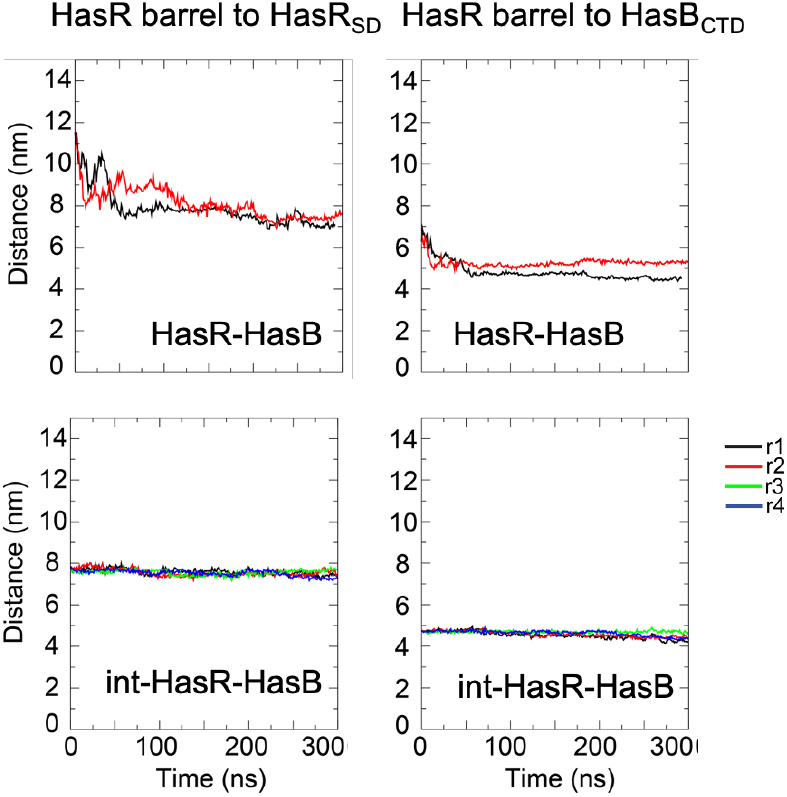
Intra- and inter-protein domain distances. The panel on the left shows the minimum distance between the HasR barrel and the HasR_SD_ as a function of simulation time. The panel on the right shows the minimum distance between the HasR barrel and the HasB_CTD_. In each case the center of mass of the protein domain is used for the measurements. Different simulations from r1 to r4 are presented.

In simulations of the int-HasR-HasB system, the major motion is a rearrangement of HasB_CTD_ with respect to the HasR linker and HasR_SD_ (since the large motion of HasR_SD_ and HasB_CTD_ had already occurred). These data strongly suggest that the HasR periplasmic region (the linker including the TonB box) is considerably flexible and that the starting model required some inter-protein adjustments in particular with respect to interactions of the HasR_SD_ with HasB_CTD_, given the adjustment occurred spontaneously during the simulations.

In order to explore these movements in a more quantitative manner, we measured the HasR barrel-HasR_SD_ and the HasR barrel-HasB_CTD_ distances. In the HasR-HasB simulations there is a reduction in the HasR barrel-HasR_SD_ distance from ∼ 11 nm in the original model to ∼7-7.5 nm by the end of both simulations, consistent with 8 nm determined in our previous work by SAXS and EM (13). Similarly, the HasR barrel to HasB_CTD_ distance is decreased from ∼ 6.8 nm to ∼ 4.5 nm in one simulation and to ∼ 5.4 nm in the other simulation. In simulations of the int-HasR-HasB system, the distances remain at ∼ 7.7 and 4.7 nm respectively across all four simulations. A striking feature of these measurements is the consistency in the int-HasR-HasB simulations. Measurements for all four independent simulations of each system stabilise to essentially the same value. This indicates that in terms of large-scale movement, the protein domains have reached a metastable state for these complexes (Figure 2).

### HasB-HasR TonB-box salt bridge

In three of the four int-HasR-HasB simulations we observed a spontaneous formation of the salt bridge between D_100_ of HasR TonB-box and R_169_ of HasB, which was previously identified by our experimental work (15) (Figure 3). Interestingly in some simulations the interaction is disrupted (defined as the minimum distance between the residues > 0.35 nm), but then reforms either for an extended period or for transitory periods.

**Figure 3:**
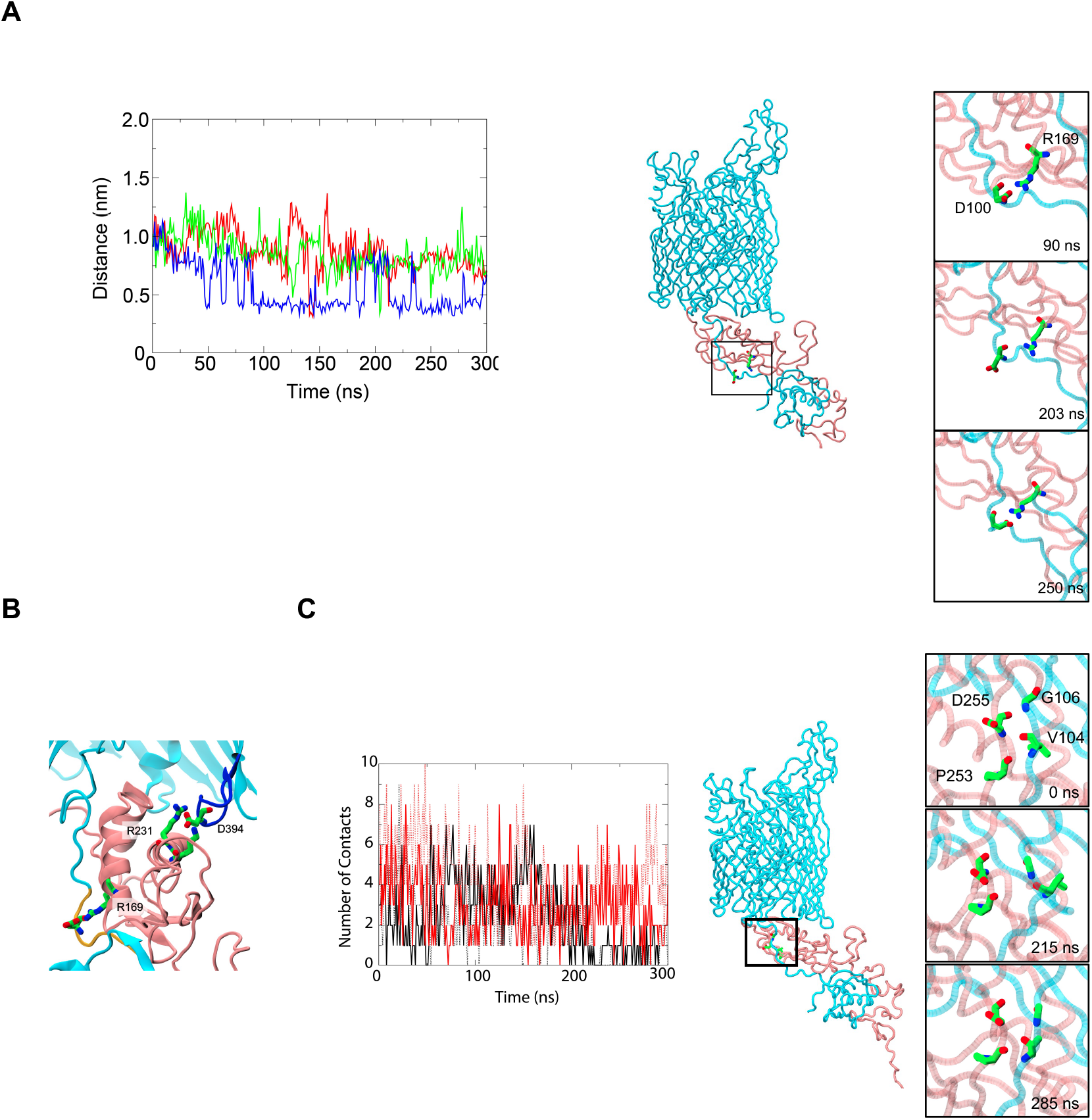
HasR TonB-box and HasB interaction. **A:** HasR TonB-box and HasB salt bridge Distance measurement between D_100_ of HasR TonB-box and R_169_ of HasB. Three simulations (in blue, red and green) are shown. **Right:** a zoomed in version of the central panel shows the two residues (D_100_ and R_169_) from one of int-HasR-HasB simulations as an example of the reformed salt bridge during simulation. **B**: HasB-HasR TonB-box salt bridge. Snapshot taken from t = 250 ns of one of the int-HasR-HasB simulations, showing the HasR D_100_-HasB R_169_ salt bridge intact. The HasR D_394_-HasB R_231_ salt bridge is also shown. HasR, L4 and the TonB box of HasR are coloured in cyan, blue and yellow respectively. HasB residues 1 to 70 and HasR residues 430 to 600 are omitted for clarity. **C:** Hydrogen bonds within the HasR-HasB inter-protein β-sheet. The panel on the left shows the total number of hydrogen bonds between HasR residues 99-111 and HasB residues 246-258 as a function of simulation time. The panel on the right (a zoomed in version of the central panel) shows four residues (V_104_ and G_106_ of HasR and P_253_ and D_255_ of HasB) highlighted from one of the int-HasR-HasB simulations as an example of the lability in the interactions in this region. At time = 0 ns, backbone hydrogen bonding is seen between HasR V_104_ and HasB D_255_; HasR G_106_ and HasB D_255_, HasR V_104_ and HasB P_253_. At 215 ns the inter-protein hydrogen bonds are broken, by 285 ns the hydrogen bond between HasR V_104_ and HasB P_253_ is reformed and the four residues are in general closer together than at 215 ns.

### HasR-HasB inter-protein β-sheet

In the initial model the residues nearby D_100_ in HasR (93-111) are located at around 3Å from the third β-strand of HasB (246-258), a distance compatible with hydrogen bonding and formation of a β-sheet, in accordance with an interprotein β-sheet as observed in our previous model between HasB_CTD_ and a peptide containing HasR TonB-box(15). We monitored the number of backbone hydrogen bonds between the two proteins in this region in each simulation (Figure 3C). There are up to 5 hydrogen bonds present between residues 93-111 of HasR and residues 246-258 of HasB_CTD_ in any one frame of HasR-HasB and int-HasR-HasB simulations. In the HasR-HasB simulations there is a reduction in the number of hydrogen bonds as the simulations proceed, such that by ∼ 200 ns there are generally either none or just a single hydrogen bond present in this region. In the int-HasR-HasB simulations there are 2-8 hydrogen bonds throughout the simulations. The number of hydrogen bonds does decrease as some of the simulations proceed. Hydrogen bonding between backbone atoms is in some cases replaced by hydrogen bonds between side chains in this region. Examples include the backbone hydrogen bonding interaction between HasB P_253_ and HasR V_104_, which is replaced by HasB P_253_ (backbone) to HasR T_103_ (side chain) in one of the int-HasR-HasB simulations. Full details are provided in the supplementary information (Figure S3). Thus, this region of interaction is labile and while the strands remain in close proximity to each other, the number of hydrogen bonds fluctuates. Overall, the inter-protein contacts are more stable in the int-HasR-HasB simulations, in other words once contact with HasR periplasmic loops has been established. We note here that an equivalent inter-protein β-sheet was reported in the X-ray structure of the complexes between *E. coli* TonB_CTD_ and the TonB-box of the TBDT FhuA, BtuB and FoxA (6–8).

### Interaction of HasR periplasmic loops with HasB

As already discussed, at the start of the HasR-HasB simulations there is a large inter-protein motion in which the distance between the HasR_SD_ and the HasB_CTD_ is reduced. In one of the simulations this movement results in interactions of HasB with HasR periplasmic loops L1 and L4, the loop numbering being from the N- to the C-terminus. The interactions formed spontaneously within about 20 ns for L1 and 50 ns for L4, and then remained stable for the duration of the simulation. The int-HasR-HasB simulations were initiated from when these loops have already made contact with HasB_CTD_, and these interactions are maintained for the duration of all four simulations (4 × 300 ns). The HasR L1-HasB interactions are largely through hydrophobic contacts of A_238_ and P_239_ of HasR L1 with L_189_ of HasB_CTD_ (Figure 4A, 4B).

**Figure 4.**
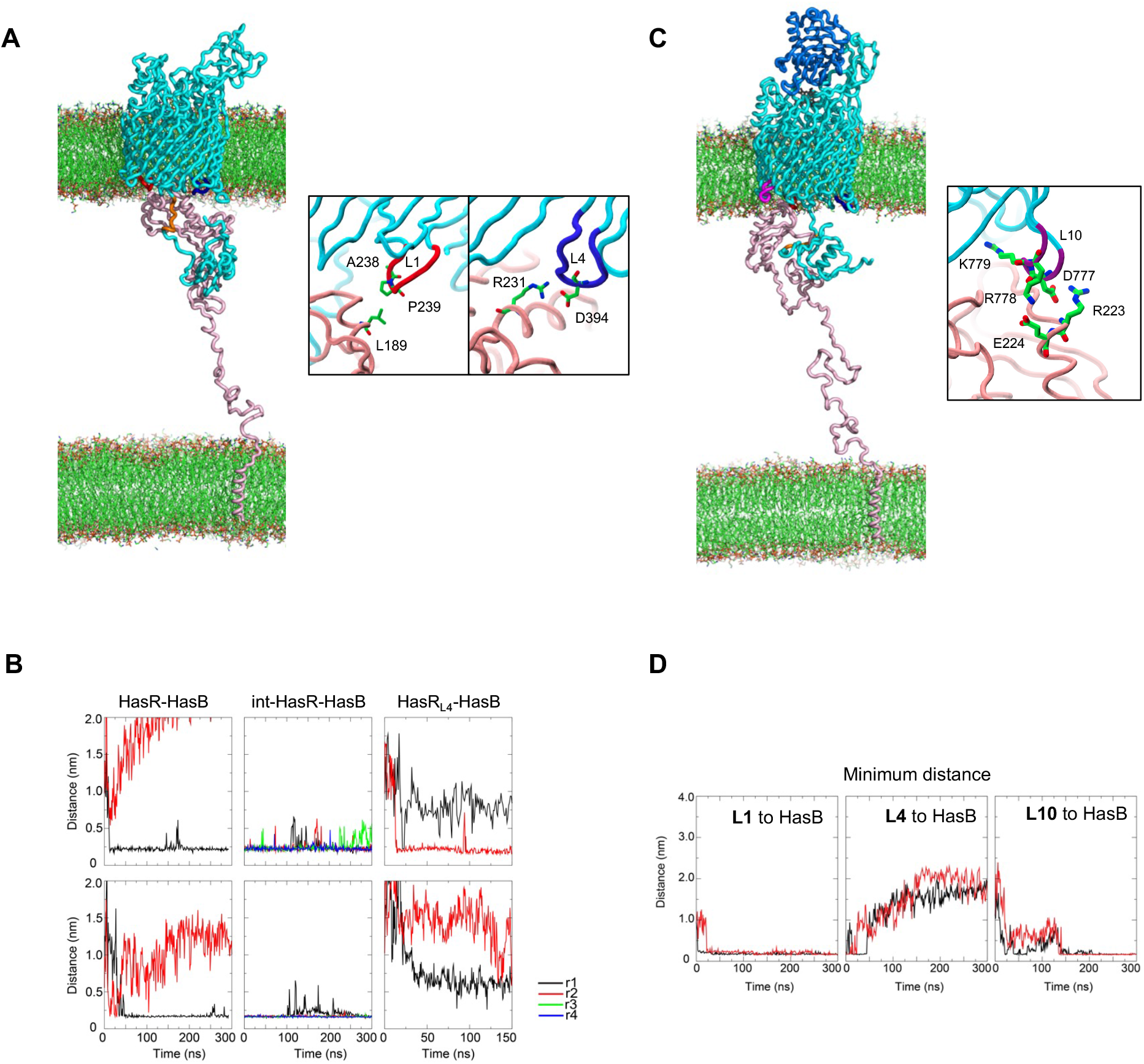
Interaction of HasR periplasmic loops with HasB. **A: HasR-HasB**. Left: Snapshot of one of the simulated HasR-HasB complex in membranes. HasR L1 and L4 are coloured in red and dark blue, respectively. The TonB box is in orange. Right: Some key residues are labelled and shown. **B**: Evolution of the minimum distance between HasR L1 (top row) or L4 (bottom row) to HasB during different HasR-HasB simulations (r1, r2, r3 and r4) are shown. The distance from the closest atoms between the loops and HasB was measured. **C: HoloHasR-HasB**. Left: Snapshot of the simulated holoHasR-HasB complex in membranes at 285 ns. Same colour codes as in A. L10 is coloured in purple. HasA in blue and heme in black. **D**: Evolution of minimum distance between HasR L1, L4 or L10 to HasB during the holoHasR-HasB simulations. Two simulations are presented, r1 in red and r2 in black.

Once the interactions with L1 are formed, the HasR linker and HasB_CTD_ move with respect to each other and then HasB_CTD_ contacts L4 *via* several salt bridges. The most prevalent are between either E_393_ or D_394_ of HasR L4 with either K_230_ or R_231_ of HasB_CTD_ (Figure 4A, 4B). Salt bridges between E_393_ and R_223_ are also observed. While the L4-HasB_CTD_ interaction exists for the duration of the 5 × 300 ns of the simulations in which it is observed, no other individual salt bridge is observed for the duration of any of these simulations. Instead, the salt bridges are observed to form, break and often reform throughout the simulations, suggesting either a protein-protein interface stabilised by non-specific interactions or that the simulations are not long enough to achieve a stabilised interface between the two proteins. Having said that, the persistence in the overall number of contacts in all five simulations between the proteins in this region does suggest that L4 is important for the interaction of HasR with the HasB_CTD._ This is further reinforced in the simulation on the HasR_L4_-HasB system, HasR_L4_ being a variant of HasR in which L4 (P_392_EDVDWLD) is replaced by the sequence ATSA. The size of four residues is the minimum required to form a periplasmic loop. In these simulations the distance between the centres of mass of the HasR_L4_ barrel and HasB_CTD_ is ∼ 5.6 nm (compared to ∼ 4.5 nm in the simulations in which contact between L4 and HasB_CTD_ is observed) and L4 does not come within hydrogen-bonding distance of HasB_CTD_ during these simulations (Figure 4B). Interestingly in the absence of L4-HasB interaction, the salt bridge between D_100_ of HasR TonB-box and R_169_ of HasB is not maintained.

### Effect of the periplasmic thickness in the HasR-HasB interaction

The HasR-HasB interaction occurs in the periplasmic space. Periplasmic thicknesses from 10 to 50 nm have been reported. Therefore, we investigated the effect of the periplasm thickness on the HasR-HasB interactions observed in our original simulations by extending the width of the periplasm to 23 nm (measuring from membrane midpoints). We constructed two models with a more extended HasB proline-rich domain to span the wide periplasm and performed two independent simulations of each, giving a total of 4 × 50 simulations. The hydrogen bonds between residues involved in the inter-protein β-sheet (Figure S4A) and the HasRL4-HasB and HasRL1-HasB (Figure S4B) interactions were maintained throughout these simulations. We also observed clustering of hydrophobic residues in the HasB proline-rich domain (Figure S4C), however these were not in the same location as those observed within the narrower periplasm. Interactions of HasB periplamic region (*i*.*e*. residues 90-256, a part of the proline-rich domain and HasB_CTD_) and POPE lipids were also observed in these simulations (Figure S2B). These data not only show that the HasR-HasB interactions are stable in the wide periplasm, but also that the HasB proline-rich domain has considerable conformational flexibility and can likely adopt a range of conformations at different steps of the mechanism.

### Effect of the presence of heme and HasA on the HasR-HasB interaction

The heme internalization through HasR and the ejection of HasA need the energy that is provided by HasB through the HasR-HasB interaction. In a previous study, we showed that the presence of heme and HasA on the extracellular side of HasR modified its interaction with HasB^14^. Here, in order to get insight into this signaling process and identify the interactions that are modified, we performed molecular dynamics simulations of the interaction of HasR loaded with HasA and heme (holoHasA) with HasB. We performed 2 × 300 ns simulations of the holoHasR-HasB system. Given the importance of the D_100_-R_169_ salt bridge for the heme internalization *via* HasR, we analysed the trajectory in which this salt bridge is most stable.

As in the apo-form, at the early stage of the simulation L4 and L1 of the holoHasR interact with HasB. Whereas the L1-HasB interaction stays stable during the simulation, the interaction with L4 is lost. Instead, the loop 10 (L10) binds to HasB. This interaction involves D_777_R_778_K_779_ of HasR L10 and R_223_E_224_ of HasB, and stays stable during the simulation (Figure 4 C, 4D).

In order to further understand the importance of L10 in this interaction, we performed molecular dynamics simulations with a variant of holoHasR in which L10 residues (R_774_AFDRKLD) were replaced by ATSA. Interestingly in this mutant, the L4-HasB interaction is formed and remains stable. Although in this mutant complex, the residues of HasB interacting with L4 are different from those in the HasR-HasB simulations presented above (R_178_, R_181_, R_188_ *versus* K_230_R_231_), the orientation of HasB with the respect of the β barrel is similar, suggesting a preferred orientation (Figure S5). During these simulations, the salt bridge between the TonB-box and HasB is stable, and the HasB-HasR barrel and HasR_SD_-barrel distances are comparable to that observed in other complexes in this study.

### Experimental validation of the role of HasR loops in the HasR-HasB interaction

In previous work, we studied a HasR mutant in which the TonB-box was deleted (HasR_ΔTBB_). Although HasR_ΔTBB_ was not functional *in vivo*, it was still able to bind to HasB_CTD_ *in vitro*, but with significantly less affinity (16) and lower ΦλH (Figure 5). We concluded that besides the TonB-box, other regions of HasR are recognized by HasB. In the present study, the simulations revealed the importance of HasR periplasmic loops L1, L4 and L10 in the HasR-HasB interaction. Here we measured the thermodynamic parameters of the interaction between HasB_CTD_ and variants of HasR in which the sequence of L1 (P_232_GKE), L4 (P_392_EDVDWLD) or L10 (R_774_AFDRKLD) were substituted by ATSA. These variants lacked the whole N-terminal region containing HasR_SD_ and the TonB-box (HasR_ΔNter_), to evaluate only the contribution of periplasmic loops in the interaction. The HasR mutants HasR_ΔNter-L1,_HasR_ΔNter-L4,_HasR_ΔNter-L10_ were prepared and purified by using the same protocol as for the wild type, indicating that they were well folded and inserted into the OM. The effect of the L4 mutation is drastic, as only a small endothermic signal that could correspond to a heat of dilution was observed in the ITC experiment with HasR_ΔNter-L4_ and HasB_CTD_ (Figure 5 top). In the simulations, HasR L4 forms polar interactions and salt bridges with K_230_R_231_ of HasB. Similarly, the ITC titration of HasB_CTDK230AR231A_ and HasR_ΔNter_ show an endothermic signal (Figure S6). The enthalpic signal reflects the contribution of polar interactions. The loss of the enthalpic signal observed with these mutants compared to the ΔH value (−7.3 kcal.mol^-1^) measured for the HasR_ΔTBB_-HasB_CTD_ interaction is compatible with at least two salt bridges that could correspond to the contribution of the HasR L4-HasB K_230_R_231_ interaction.

**Figure 5.**
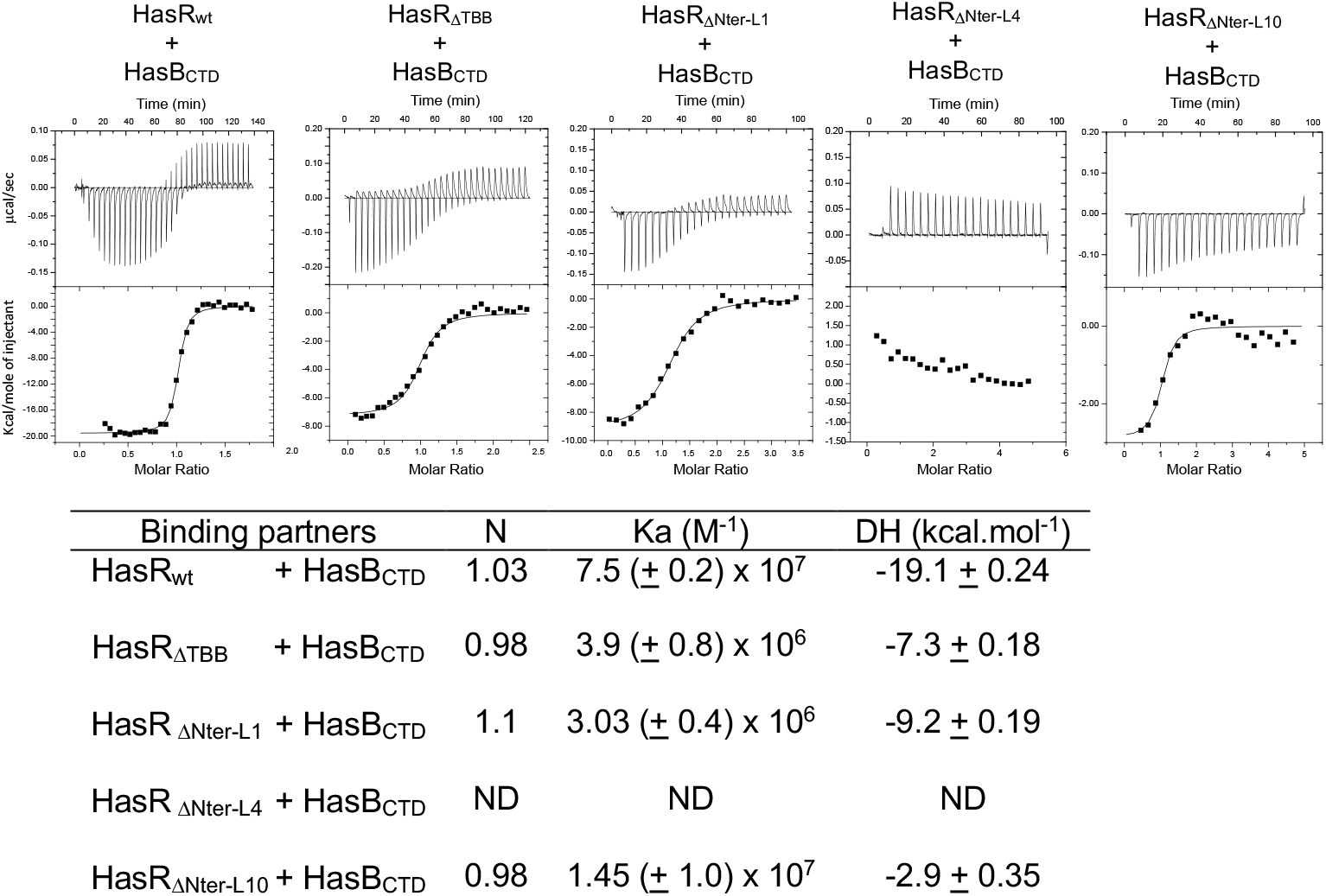
ITC analysis of the interaction of HasB_CTD_ with different variants of HasR. Top: Representative experiments are shown. In each case, the heat signal is shown together with the binding isotherm derived from the signal. Bottom: Stoichiometries (N), affinity constants (Ka) and ΔH of binding at 25°C are shown. ND: not determined.

HasR_ΔNter-L10_ still interacts with HasB_CTD_, although with less affinity (3.7 times) and with a loss of polar interactions compared to HasR_!’1TBB_ (ΦλH -2.9 *versus* -7.3 kcal.mol^-1^) (Figure 5). In the simulations, HasR L10 residues form salt bridge and polar interactions with HasB R_223_E_224_. Unfortunately, the HasB_CTD_ variant in which these charged residues are replaced by neutral ones, oligomerises and is not stable as monomer. Therefore, we could not study the interaction of this mutant with HasR *in vitro* by ITC.

HasR_ΔNter-L1_ interacts with HasB_CTD_ with comparable thermodynamic parameters as HasRΔTBB showing that the substitution in L1 has no effect on the binding and on the contribution of polar interactions. This is consistent with the simulations showing that this loop interacts with HasB_CTD_ by mostly hydrophobic contacts.

To assess the importance of HasR L4 and L10 in the mechanism of heme entry, we monitored the growth of *E. coli* strain *K12* C600*ΔhemAΔtonB* that needed an external source of heme (here 0.4µM heme-BSA) and expressed the complete <u>has</u> locus (*hasISRADEB*) with either *hasR* wild type or mutated in L4 (HasR_L4_) or in L10 (HasR_L10_). A mutant in L3 (HasR_L3_) was used as a control. The complete *has* allows the expression of all Has proteins required for the secretion of HasA (*hasA, hasD, hasE*), heme internalisation (*hasR, hasB*) and the transcription regulation of the system (*hasI, hasS*). In this *E. coli* strain *tonBexbBexbD* were removed and *exbBexbD* from *S. marcescens* was added (HasB partners in the IM) in order to have a fully functional HasB.

The growth curves with the expression of the HasR wild type, HasR_L3_, HasR_L4_ or HasR_L10_ are shown in Figure 6. The mutation on HasR L4 and L10 significantly affect the bacterial growth curve (Figure 6A). HasR_L3_ is used as a control. The expression level of all variants is comparable to that of the wild type, indicating that the mutations do not modify the overall folding of the protein (Figure 6A). The lag phase is significantly longer with HasR_L4_ (9.5h) and HasR_L10_ (25h) versus 5.5h for the wild type and HasR_L3_. In addition, with the two variants HasR_L4_ and HasR_L10_, the OD600 do not reach the maximum obtained with the wild type HasR. The most important effect was observed with HasR_L10_ where only an OD600 at 0.1 is reached after 40h of growth, compared to the wild type with an OD600 at 0.57 after 25h. These observations are consistent with the key role of L4 and L10 in the interaction with HasB revealed by our simulations and observed in ITC experiments.

**Figure 6.**
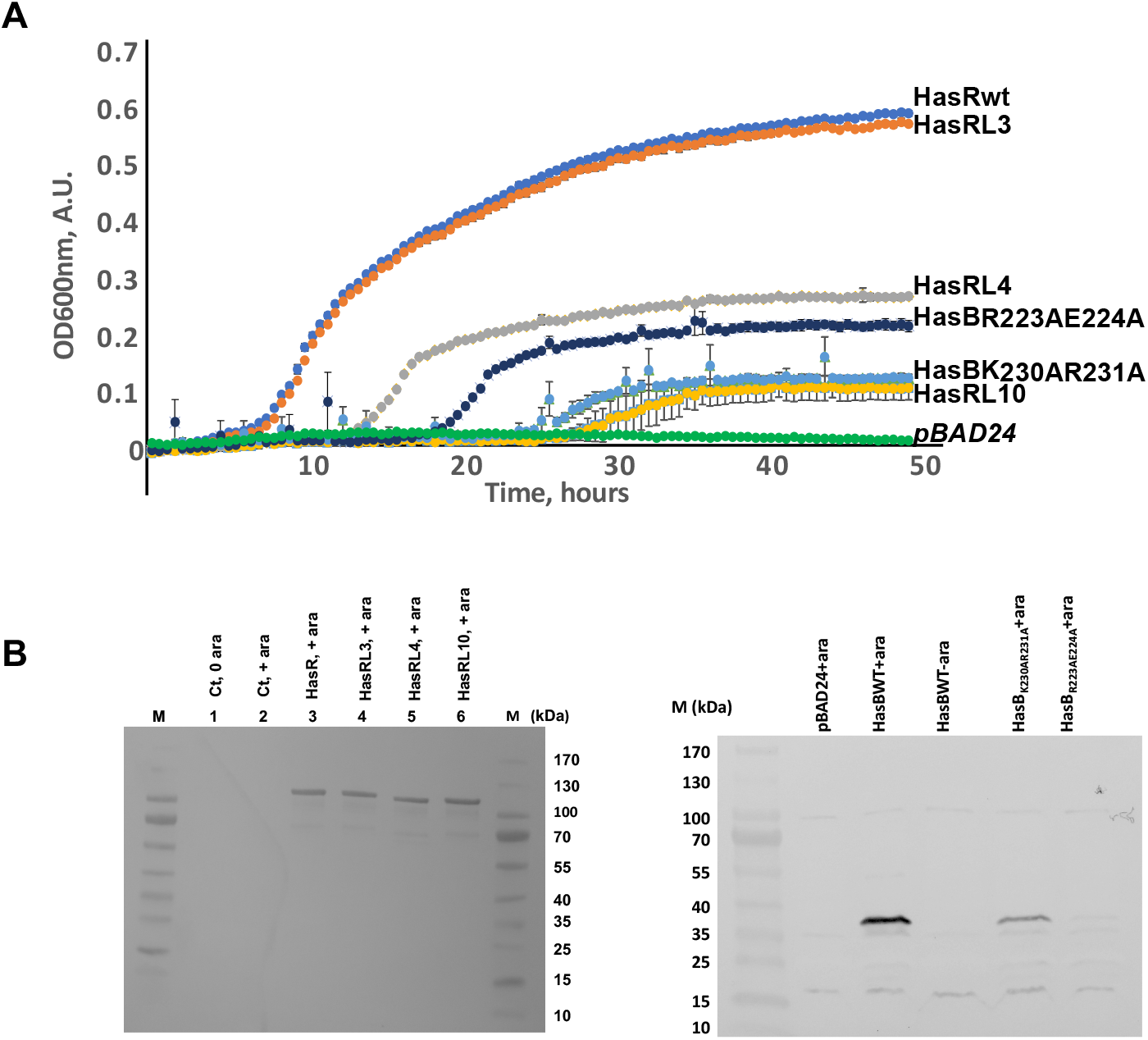
Phenotypic analysis of HasR periplasmic loops substitution in vivo. **A:** Growth curves of E. coli C600 ΔhemAΔtonBΔexbBD harbouring pBAD24 vector (negative control) or its derivatives encoding exbBD_sm_ and the wild type has locus (HasR wt), HasR derivatives with HasR L3, L4 or L10 substitution or HasB derivatives are presented and labelled. **B:** Top, Immunodetection with polyclonal anti-HasR rabbit antibody in C600ΔhemA transformed with either pBAD24 (negative control, line 1 and 2), pFR2 (HasR_wt_, line 3), pFR2: HasR_L3_, pFR2: HasR_L4_ or pFR2: HasR_L10_. Cells were inoculated at 0.05 OD_600nm_ in LB medium containing δ-aminolevulinic acid (25 µg/ml) at 37°C for 3 hours. An equivalent of 0.1 OD600nm was loaded on each lane. Arabinose (Ara) concentration was 40µg/ml. Molecular mass of standard weight markers is indicated on the right. Bottom: Expression level of HasB, HasB_K230AR231A_, HasB_R223AE224A_ measured by immunodetection with polyclonal anti-HasB rabbit antibody in the E. coli K12 strain JP313 (ΔaraBAD) was used and transformed with a plasmid expressing ExbBD from S. marcescens and another plasmid expressing HasB or its mutants or a control plasmid (pBAD24). Induction was with 40µg/ml arabinose for two hours at 37°C. The equivalent of 0.15 OD_600nm_ was loaded on each lane. After transfer on a nitrocellulose membrane and incubation with the primary antibody (1/5000 dilution), the immunoblots were revealed with goat anti-rabbit IgG coupled to alkaline phosphatase.

In the simulations, L4 forms stable polar interactions and salt bridges with K_230_R_231_ of HasB, and L10 with R_223_E_224_. We substituted these charged residues to alanine and tested the activity of the mutants HasB_K230AR231A_ and HasB_R223AE224A_ *in vivo* (Figure 6A). These mutants give comparable phenotype than the HasR_L4_ and HasR_L10_ variants, although the level of HasB_R223AE224A_ expression and/or stability is lower than that of the wild type (Figure 6B).

Together the *in vivo* results confirm the key role of HasR L4 and L10 in the interaction with HasB_CTD_ and in the HasR function.

## Discussion

Our study reveals the key role of specific periplasmic loops of a TBDT in its interaction with a TonB protein and in its function. Moreover, the simulations show that the involved loops and the sequence of the events are different depending on the presence of the extracellular substrates, heme and HasA. This implies that the conformational and dynamics changes associated to substrate binding on the extracellular side of HasR are transmitted through the barrel and plug to the periplasmic side of HasR and the region of interaction with HasB.

In the absence of extracellular substrates, the interaction of HasR L1 and HasB occurs very early, mainly *via* hydrophobic contacts. Once these contacts are formed, the HasR linker and HasB move with respect to each other (mostly through HasB rotation) such that HasB contacts L4 (Movie S1). The HasR-HasB interaction is then stabilized through several salt bridges between L4 and HasB K_230_R_231_ residues (Figure 4A). When the charged residues involved in this interaction were replaced by neutral residues, either in HasB (HasB_K230AR231A_) or in HasR (HasR_L4_) the bacterial growth is significantly affected (Figure 6), suggesting the key role of the L4-HasB interaction in heme import and/or signalling through HasR. The ITC experiments showed also that L4 is required for HasR-HasB complex formation and stability (Figure 5). The HasR L4-HasB interaction seems to stabilise the salt bridge between the HasR TonB box and HasB, since in the simulations with HasR_L4_, this salt bridge is not maintained. As shown previously, this salt bridge is required for HasR function *in vivo* (16).

When the extracellular substrates are present on HasR (holoHasR), although L4 and L1 form interactions with HasB quickly, the interactions with L4 are disrupted while those with L1 are maintained. Instead, L10 residues form polar and ionic interactions with HasB (Figure 4C). We validated the key role of L10 and of its charged residues in the interaction with HasB *in vitro* and with functional *in vivo* assays (Figure 5 & 6). Interestingly in the simulation where the charged residues of L10 are substituted by neutral ones (holoHasR_L10_-HasB), instead of L10-HasB, the L4-HasB interaction is formed (Figure S5). This is consistent with the *in vivo* data showing that the substitution in L10 or L4 does not completely abolish the heme acquisition and suggests that L4 and L10 can be involved in different steps of the heme acquisition and might be partially interchangeable. In our simulations, the L10-HasB interaction occurs only when HasR is in its holo-form. This interaction could be crucial for the step of heme internalisation and the ejection of HasA, two steps for which energy is required.

Structural data showed that the TonB-TBDT interaction involves a salt bridge between the TonB-box of the transporter and TonB, stabilizing an inter-protein β-sheet (4),(6),(7),(15). Both salt bridge and the inter-protein β-sheet were shown important for the energy transfer from the TonB protein to the transporter (20),(21). In all the simulations presented here, the salt bridge between HasR and HasB and the inter-protein β-sheet are formed, although sometimes they are observed to disrupt and reform. They are more stabilised once HasB contact HasR periplasmic loops (Figure 3).

Simulations of HasR in an OM model containing lipopolysaccharide (LPS) at the Ra-LPS level (*i*.*e*. without the O-antigen group) showed very little motion of the protein in 500 ns (Figure S7). This is not surprising given the slow-moving nature of LPS. While we and others have reported that the conformational behaviour of TBDTs can be altered by the presence of LPS (22, 23), the large size of the systems we have studied here, would require simulations considerably beyond the scope of the present study to explore any impact of LPS.

If we compare the HasB-HasR interaction network with the sequence and the structure of other available TBDT-TonB complexes (PDB entries 2GSK, 2GRX and 6I97), the residues of HasR L1, L4 - notably the charged residues of L4- are well conserved in FoxA, BtuB and FhuA, although the loops are slightly shorter in these proteins compared to those of HasR. L10 is less conserved (Figure S8). The HasB residues that contact L1, L4 and L10 in our simulations are also well conserved and exposed at the surface of *E. coli* and *P. aeruginosa* TonB (Figure 7). However, no simultaneous interactions between TBDT L1 and L4/L10 with TonB are observed in these complex structures. This may be a result of only the globular C-terminal part of TonB being used in these studies. In our full-length model of HasB inserted into the IM, the dynamics of its periplasmic domain attached to the IM allows a more thorough exploration of the exposed surface of HasR and the simultaneous interaction of HasB with L1 and L4 or L10. Therefore, we hypothesise that the HasR-HasB interacting mode revealed in this study can occur in other TBDT-TonB interactions.

**Figure 7.**
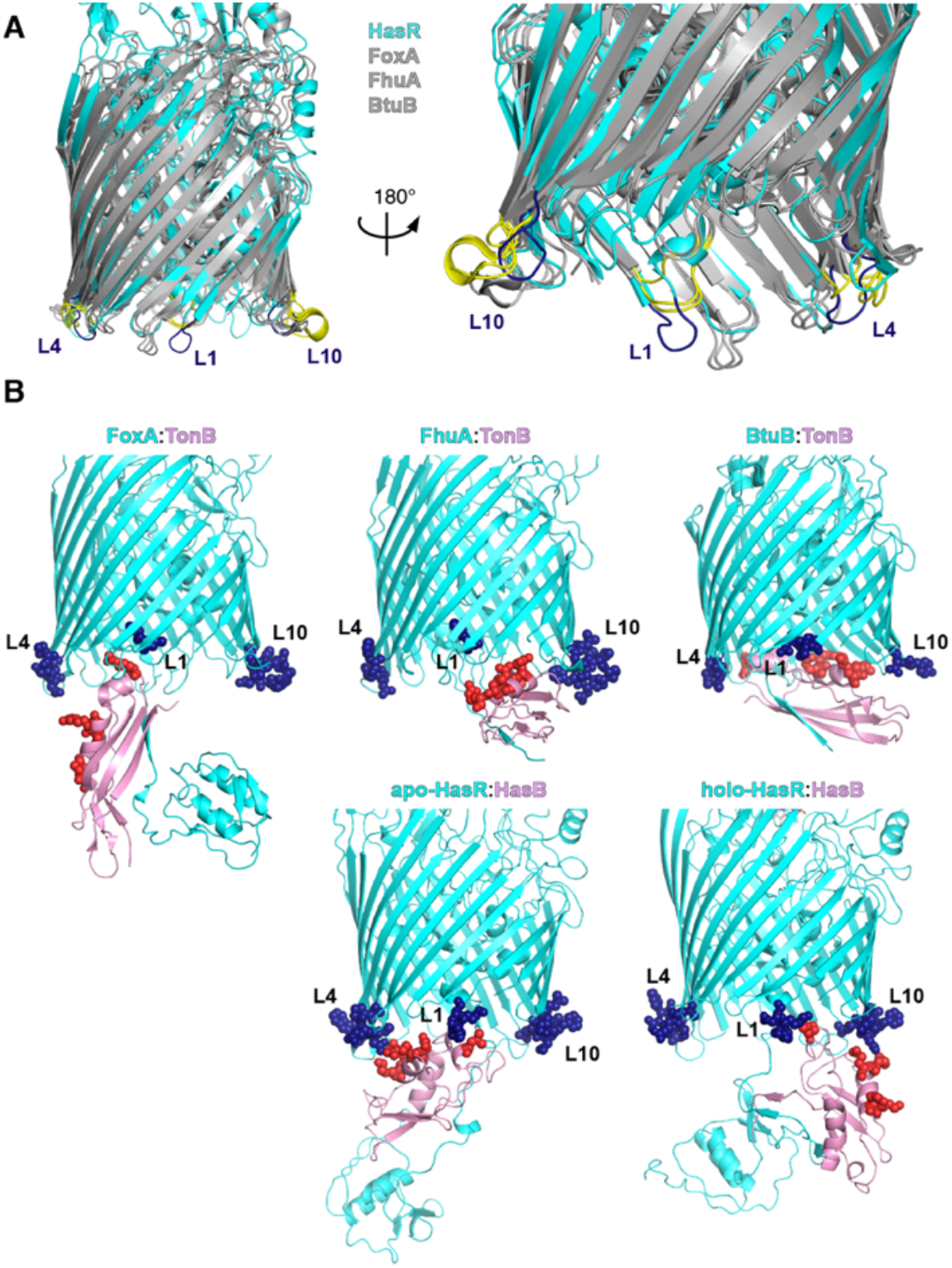
Periplasmic loops in TBDT in complex with TonB. **(A)** Superposition of TBDT structures HasR (cyan), FoxA, FhuA and BtuB (grey). Loops L1, L4 and L10 of HasR are coloured in dark blue while the corresponding residues in FoxA, FhuA and BtuB are coloured in yellow. **(B)** Transporters (cyan) in complex with the C-terminal domain of TonB/HasB (pink) are depicted in cartoon. The residues of loops L1, L4 and L10 are shown in blue spheres. Residues of TonB equivalent to the ones from HasB interacting with HasR in MD simulations (i.e. HasB L_189_, R_223_, E_224_, K_230_ and R_231_) are shown as red spheres.

The energizing mechanism used by Gram-negative bacteria to internalize scarce and essential nutrients is still not well deciphered. Understanding it will help the future development of new antibacterial strategies. It is assumed that energized TonB applies a mechanical force to remodel the TBDT plug and open a channel through the barrel, allowing substrate entry. How TonB applies this force is not yet understood. EPR (24) and AFM (21) data, as well as steered molecular dynamics (20) support a pulling movement of TonB, while a recent structural study suggests a rotational model based on structural homology to the Mot flagellar system (9). During our simulations, we observed the rotation of HasB with respect to HasR, while the HasB_CTD_-HasR barrel distance remains stable (4-5 nm in all the simulations). These observations may suggest a rotational model, rather than a pulling model for which a variation of the HasB_CTD_-HasR barrel distance would have been expected. However, the simulations are performed in the absence of ExbB and ExbD, the partners of HasB in the IM, and without the input of energy. Further studies with ExbB and ExbD will thus be required to fully understand the mechanism of energy transfer between the two bacterial membranes.

## Materials and Methods

### Positioning the protein complex in the membranes

The CHARMM-GUI web-server (25),(26) was used to assemble the lipid bilayers, both as squares with side-length 16.5 nm. The inner membrane was made up of two leaflets of 1-pamitoyl-2-oleoyl-sn-glycero-3-phosphoethanolamine (POPE). The outer membrane similarly had a lower leaflet of POPE but an upper leaflet of 1,2-dilauroyl-sn-glycero-3-phosphocholine (DLPC) as the tail lengths of DLPC are similar to those of lipopolysaccharide (which is found in the outer leaflet *in vivo*, but requires long simulations to achieve convergence and thus was not used here as our interest was in the protein complex rather than specific details of protein-outer membrane interactions)(27). The two membranes were subjected to a 5000-step energy minimisation using the steepest descents algorithm.

To align the HasR barrel in the outer membrane, co-ordinates were taken from the MemProtMD database (28) and used for alignment to ensure the lipid headgroups were in the correct position relative to the protein. The location of the HasB helix in the inner membrane was estimated by using the location of the tryptophan residues, given there was no structural information available for this section (29).

Simulations involving HasR and HasB were set up as follows; once the proteins had been inserted into the membranes, any overlapping lipids were removed prior to energy minimisation. The CHARMM-GUI webserver was used to generate the necessary topology and parameter files for all models and structures reported in this study.

### Molecular Dynamics Simulations

The systems were solvated with the TIP3P water model. K^+^ and Cl^-^ ions were added to achieve an overall concentration of 150 mM in charge neutral systems. The systems were then subjected to a short energy minimisation of 5000 steps, again by using the steepest descent algorithm, to remove any steric clashes, followed by an equilibration protocol as shown in Table S1. This protocol, with slow relaxing of the restraints on the protein was necessary to allow the lipids to equilibrate around the proteins.

All simulations were carried out with the GROMACS 2019.6 (30) version (www.gromacs.org) and CHARMM36 (31) forcefield. The long-range electrostatic interactions were treated by using the Particle Mesh Ewald (PME)(32) method, whereas the nonbonded interactions and the short-range electrostatics were cut off at 1.2 nm, applied by using the potential shift Verlet cut off scheme. The temperature was maintained at 303.15 K by using the Nosé-Hoover thermostat (33)(34) with a 5 ps coupling constant. To maintain a constant pressure at 1 atm, the Parrinello-Rahman barostat (35) was applied in a semi-isotropic fashion, again with a coupling constant of 5ps. The LINCS (36) algorithm was applied to all atoms to allow the time step of 2 fs. Table 1 shows the summary of production runs, all of which were run in the NPT ensemble.

**Table 1:** Summary of the Equilibrium MD Simulation Systems

| System | Temperature (K) | Simulation Length (ns) |
| --- | --- | --- |
| HasR + HasB | 303.15 | 300 (x2) |
| *Int-HasR + HasB | 303.15 | 300 (x4) |
| holoHasR + HasB | 303.15 | 300 (x2) |
| HasR<sub>L4</sub>+ HasB | 303.15 | 150 (x2) |
| HasR<sub>L10</sub>+ HasB | 303.15 | 150 (x2) |
| **Int-HasR + HasB in the wide periplasm | 303.15 | 50 (x2) |
| ***HasR + LPS | 303.15 | 500 (x2) |

\*The int HasR\_HasB simulations were initiated from a structure extracted after 100 ns from one of the HasR + HasB simulations after the contact with L1 and L4 had already been established.
\*\*The Int-HasR + HasB in the wide periplasm simulations were initiated from the same structure as the int HasR\_HasB simulations, but the inner membrane with HasB helix was translated to be lower ~ 8 nm. The proline-rich domain from residues 37 to 140 was remodelled by using Modeller software. To allow the linker region to improve its structure, the entire protein structure, except the modelled linker region, was subjected to position restraints of 1000 kJ mol<sup>-1</sup>. nm<sup>-2</sup> for 50 ns using gromos96 54a7 force field (37). The equilibrated protein model was then placed in the double membrane model, followed by a short equilibration with a position restraint of 50 kJ mol<sup>-1</sup> nm<sup>-2</sup> on the backbone for 10 ns by using CHARMM36 forcefield.
\*\*\*The HasR + LPS simulations were built by embedding the HasR barrel and plug domain (PDB entry 3CSL) in the outer membrane with the predefined LPS model of *E. coli* in CHARMM-GUI. The lower leaflet of the outer membrane contains POPE, 1-palmitoyl 2-cis-vaccenic phosphatidylglycerol (POPG) and tetraoleoyl cardiolipin (TOCL2) in the ratio 18:1:1 (38)(39), while the upper leaflet of the outer membrane contains a predefined Ra-LPS model of *E. coli* available in CHARMM-GUI (26, 27). The HasR with the Ra-LPS model was then subjected to the equilibration steps shown in the Table S1.

### Isothermal Titration Calorimetry (ITC)

ITC experiments were performed at 25°C by using a MicroCal VP titration calorimeter (MicroCal-GE Healthcare) under the same conditions as described previously (16). The buffer was 20 mM sodium phosphate, pH 7, 0.08% Zwittergent 3–14 for all titrations. All protein samples were dialyzed in this buffer before the experiments. Consecutive aliquots of 5-10 μL of HasB_CTD_ at 80-120 μM were added into the ITC cell containing HasR wild type or mutant proteins at 4-15 μM. The heat of dilution of protein injections was determined either by injecting the ligand into the buffer alone or by injecting the ligand into the cell after the saturation of the protein binding site. The value obtained was subtracted from the heat of reaction to give the effective heat of binding. The resulting titration data were analysed by using the Microcal-ORIGIN software package. The molar binding stoichiometry (N), association constant (Ka; Kd =1/Ka) and enthalpy changes (ΔH) of binding were determined by fitting the binding isotherm to a model with one set of sites. All experiments were done in duplicate.

### Bacterial growth test

Growth tests in liquid medium were performed with *E. coli* C600 *ΔhemAΔtonBΔexbBD* strain harbouring vector pAMhasISRADEB encoding the whole <u>has</u> locus (*hasISRADEB)* and including either the wild type *hasr* or a mutant. The expression of this vector is under Fur regulation, induced in iron deficiency conditions. An additional *vector* pBAD24exbBDsm encoding *E*xbB and ExbD from *S. marcescens*, under the control of the P_*araBAD*_ promoter was also present. A few colonies of this strain were first inoculated in 4ml of LB medium at 37°C with 100µM dipyridyl (an iron chelator), 4µg/ml arabinose and the corresponding antibiotics. Once the culture reached an OD_600nm_ between 1.2-1.5, it was diluted and inoculated in 48 well Greiner plates, in the same medium to which was added 0.4µM He-BSA, as a heme source. The initial OD_600nm_ of the cultures was 0.001. Each well contained 300µl of growth medium. Duplicates of each strain were made, and the plate was incubated at 37°C with vigorous shaking (500rpm) in a Clariostar Plus Microplate reader. OD_600nm_ was recorded every 30 minutes for 70 hours. All experiments were performed in triplicate. In the expression test of HasR and his mutants, δ-aminolevulinic acid, a heme synthesis precursor was added to the culture at 25µg/ml.

## Acknowledgments

This work was funded by the French Agence Nationale de la Recherche ANR HemeStockExchange, the Fondation pour la Recherche Médicale (Equipe FRM 2017M.DEQ20170839114). K.S is funded by the Development and Promotion of Science and Technology Talents Project (DPST), Thailand. SK is funded by the EPSRC grant number EP/R029407. The PD and AP lab is supported by LABEX Dynamo (LABEX DYNAMO ANR-11-LABEX-0011-01). We thank Christophe Thomas from the platform of production and purification of recombinant proteins at the Institut Pasteur for bacterial cell production in fermentor, the platform of molecular biophysics at the Institut Pasteur for providing access to the microcalorimeter and Loïc Hellio for HasB sample preparation. We thank Riccardo Pellarin for helpful discussions. The authors acknowledge the use of the IRIDIS High Performance Computing Facility, and associated IT support services at the University of Southampton.

